# Interplay between UNG and AID governs intratumoral heterogeneity in mature B cell lymphoma

**DOI:** 10.1101/2020.06.29.177303

**Authors:** Pilar Delgado, Ángel F Álvarez-Prado, Ester Marina-Zárate, Isora V Sernandez, Sonia M Mur, Jorge de la Barrera, Fátima Sanchez-Cabo, Marta Cañamero, Antonio de Molina, Laura Belver, Virginia G de Yebenes, Almudena R Ramiro

## Abstract

Most B cell lymphomas originate from B cells that have germinal center (GC) experience and bear chromosome translocations and numerous point mutations. GCs B cells remodel their immunoglobulin (Ig) genes by somatic hypermutation (SHM) and class switch recombination (CSR) in their immunoglobulin (Ig) genes. Activation Induced Deaminase (AID) initiates CSR and SHM by generating U:G mismatches on Ig DNA that can then be processed by Uracyl-N-glycosylase (UNG). AID promotes collateral damage in the form of chromosome translocations and off-target SHM, however, the exact contribution of AID activity to lymphoma generation and progression is not completely understood. Here we show using a conditional knock-in strategy that AID supra-activity alone is not sufficient to generate B cell transformation. In contrast, in the absence of UNG, AID supra-expression increases SHM and promotes lymphoma. Whole exome sequencing revealed that AID heavily contributes to lymphoma SHM, promoting subclonal variability and a wider range of oncogenic variants. Thus, our data provide direct evidence that UNG is a brake to AID-induced intratumoral heterogeneity and evolution of B cell lymphoma.

---

Non-Hodgkin-lymphomas (NHL) are high incidence hematological malignancies, with more than 500,000 new cases diagnosed in 2018 worldwide (World Cancer Research Fund, www.wcrf.org). NHL comprise a number of distinct pathological entities, however, the vast majority of them arise from the malignant transformation of mature B cells with germinal center (GC) experience. Those include follicular lymphoma (FL) and the more aggressive diffuse large B cell lymphoma (DLBCL), which together account for more than 50% of all NHL, as well as Burkitt lymphoma (BL) ^1^.

The etiological relevance of GCs in the malignant transformation of B cells is complex and multi-layered ^2,3^. GCs are microanatomical structures that are formed in secondary lymphoid organs upon B cell stimulation during the immune response. When activated by antigen in the context of an antigen presenting cell and a cognate T cell, B cells engage in an intense proliferative reaction and are subject to point mutations at the variable, antigen recognition portion of their antibody (or immunoglobulin, Ig) genes, a process called somatic hypermutation (SHM). SHM, combined with affinity maturation, allows the generation of higher affinity antibodies. In addition, GC are the site for a region-specific recombination reaction called class switch recombination (CSR), which replaces the primary IgM/IgD constant region by alternative downstream constant regions at the Ig heavy (IgH) locus, thus giving rise to IgG, IgE or IgA isotypes and adding enormous functional versatility to antibody-mediated antigen clearance ^4–7^.

CSR and SHM are triggered by a single enzyme called activation induced deaminase (AID) through an initial cytosine to uracil deamination event on the DNA of Ig genes. AID deficiency dramatically blocks SHM and CSR and causes human HIGM2 immunodeficiency ^8,9^. AID-mediated deamination results in a U:G mismatch that can then be recognized by base excision repair (BER) or mismatch repair (MMR) pathways to ultimately give rise to a mutation (in the case of SHM) or a double strand break and a recombination reaction, in CSR. Accordingly, Ung^-/-^-BER deficient-mice, and Msh2^-/-^ -MMR deficient-mice show different alterations in SHM and CSR ^10–17^. Likewise, double Ung^-/-^Msh2^-/-^ deficient mice only retain transition mutations at C/G, directly emerging from the replication of U:G mismatches, and are devoid of other SHM footprints, as well as of CSR ^18,19^.

Early studies showed that several lymphoma-associated oncogenes bore mutations with the hallmark of SHM ^20–22^, and the molecular contribution of AID activity to off-target SHM was directly shown in mice later ^19,23^, indicating that a relatively high proportion of genes can be targets of AID mutagenic activity. Moreover, AID can trigger chromosome translocations involving IgH and *Myc*, the hallmark of Burkitt lymphoma ^24–26^, as well as many other genomic locations ^27,28^. Notably, AID-dependent myc/IgH translocations are abolished in the absence of UNG ^24^, suggesting that UNG activity is required for the processing of AID-deamination into the DNA double strand break preceding the chromosome translocation joining. On the other hand, SHM frequency is increased in the combined absence of UNG and MSH2, indicating that BER and MMR contribute together to the faithful repair of AID-induced deaminations ^4,19,23,29^. In line with these observations, UNG deficient mice develop B cell lymphomas ^30^.

Numerous studies have correlated AID activity with human B cell neoplasia. AID expression is associated to bad prognosis in hematological malignancies ^31–35^ and footprints of SHM have been found in human B cells ^20,21,36^ and B cell neoplasia ^22,38,39^. In addition, genetic mouse models have been used to explore the causative role for AID in B cell transformation. Several loss of function approaches have shown that AID deficiency delayed the appearance of B cell malignancies ^25,33,40^ and that this effect might be specific of mature B cell neoplasias ^33,41,42 43^. However, gain-of-function approaches uncovered an unexpected complexity on the contribution of AID to B cell malignant transformation. In an early study where AID was expressed as an ubiquitous transgene, T, but not B cell lymphomas were generated ^44^; this idea of tissue- or stage-specific sensitivity to AID-induced transformation was also proposed in other transgenic models ^45^. Two additional AID transgenic mouse models failed to promote B cell transformation, presumably due to B cell specific negative regulation of AID levels ^46^ or activity ^47^. Finally, an AID transgene driven by Ig light chain regulatory regions revealed that although AID promoted genomic instability, B cell transformation required concomitant loss of p53 tumor suppressor ^48^. Thus, AID’s contribution, and specifically of SHM, to B cell transformation is not fully understood.

Here, using a non-transgenic conditional mouse model for AID supra-expression, we found that AID expression alone was not sufficient to promote B cell lymphomagenesis, regardless of the onset of AID expression and even in the absence of p53. In contrast, removal of BER activity by breeding to Ung^-/-^ mice resulted in an increase of SHM and of B cell lymphoma incidence. Molecular characterization of Ung^-/-^ tumors by whole exome sequencing revealed that AID supra-expression increased intratumor heterogeneity (ITH), giving rise to a more diverse range of oncogenic alterations. Thus, our work establishes formal genetic evidence for the contribution of AID-mediated SHM to B cell lymphoma tumor evolution.

## RESULTS

### A mouse model to address the contribution of AID to B cell lymphomagenesis

To address the contribution of AID activity to B cell lymphomagenesis we made use of a conditional mouse model for AID expression ^49^. A cassette containing the *Aicda* and *EGFP* cDNAs separated by an internal ribosomal entry site (IRES) was placed into the endogenous *ROSA26* locus preceded by a floxed transcriptional stop sequence (R26^+/AID^ mice, Figure S1A, ^49^). Thus, AID expression from the Rosa26-AID allele is silent until the transcriptional stop is removed by Cre. To approach the role of AID in B cell lymphomagenesis we induced AID expression in B cells by breeding R26^+/AID^ mice to Cd19^+/cre^ mice, where Cre is inserted in the B-cell specific CD19 allele ^50^ (R26^+/AID^ Cd19^+/cre^ mice). We observed that the onset of expression from the Rosa26-AID allele occurred early in B cell precursors and progressively increased during B cell differentiation, being almost complete in mature B cells, and consistently with previous reports using the Cd19-Cre strain (Figure S1B) ^51,52^. Expression of AID at early stages did not detectably affect bone marrow differentiation (Figure S1C).

To assess AID expression in mature B cells from R26^+/AID^ Cd19^+/cre^ mice, we performed a qRT-PCR analysis that allowed quantification of AID mRNA expressed specifically from the Rosa26-AID allele, or total AID mRNA, resulting from the combined expression from the endogenous AID allele and the Rosa26-AID allele. Expectedly, we detected expression of Rosa26-AID both in naïve and activated B cells from Rosa26^+/AID^ Cd19^+/cre^ mice and not in B cells from control mice (Figure 1A, top graphs). Accordingly, assessment of total AID mRNA showed that the Rosa26-AID allele promotes expression of AID in naïve B cells (Figure 1A, bottom left) and significantly increases the total expression level of AID in activated B cells (Figure 1A, bottom right). In agreement with this result, western blot analysis in naïve B cells revealed AID protein only in R26^+/AID^ Cd19^+/cre^ mice, while LPS+IL4 activated B cells had increased AID protein levels in R26^+/AID^ Cd19^+/cre^ mice, compared to R26^+/+^ Cd19^+/cre^ control mice (Figure 1B). Thus, we have generated a model for early expression of AID in naïve B cells and supra-expression in activated B cells.

**Figure 1.**
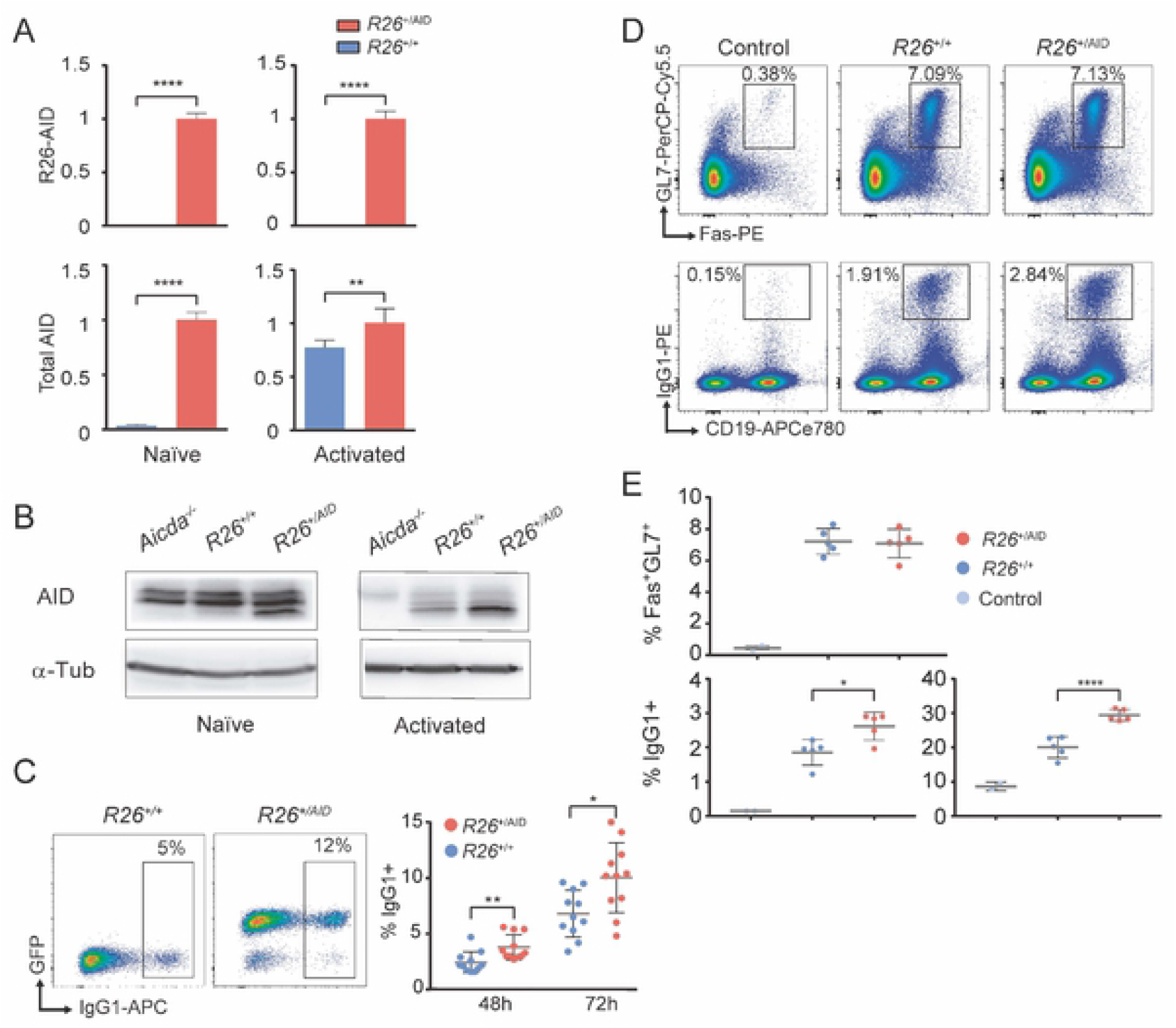
Characterization of R26^+/AID^ Cd19^+/Cre^ mice. **(A)** Increased expression of AID mRNA in B cells from R26^+/AID^Cd19^+/Cre^ mice. mRNA levels of R26-AID and total AID was measured by RT-qPCR in naïve (left) and in LPS+IL4 activated B (right) cells from R26^+/AID^Cd19^+/Cre^ (red bars) and R26^+/+^Cd19^+/Cre^ control mice (blue bars). Values are relative to levels in naïve (left) or activated (right) B cells from R26^+/AID^Cd19^+/Cre^ mice; mean of two experiments is shown; n=6 per group). **(B)** Increased AID protein in B cells from R26^+/AID^Cd19^+/Cre^ mice. Total lysates of naïve or LPS+IL4 activated B cells from R26^+/AID^Cd19^+/Cre^ and R26^+/+^Cd19^+/Cre^ mice were analysed by western blot with a monoclonal anti-AID antibody. Incubation with an anti-tubulin antibody is shown as loading control. **(C)** Increased CSR in R26^+/AID^Cd19^+/Cre^ mice. Representative FACS analysis of CSR to IgG1 in LPS+IL4 activated B cells from R26^+/AID^Cd19^+/Cre^ and R26^+/+^Cd19^+/Cre^ mice (left panel). Quantification is shown on the right. Each dot represents an individual mouse (n=11mice per group). **(D)** Increased immunization response in R26^+/AID^Cd19^+/Cre^ mice. Representative FACS analysis of GC (Fas+GL7+) B cells (upper panels) and CSR to IgG1 (lower panels) in non-immunized R26^+/+^Cd19^+/Cre^ (control, PBS) and SRBC-immunized R26^+/+^Cd19^+/Cre^ and R26^+/AID^Cd19^+/Cre^ mice after B220+ gating. **(E)** Quantification of Fas+GL7+ GC B cells (upper panel) and CSR to IgG1 (lower graphs) on B220+-gated (lower left) and B220+Fas+GL7+-gated B cells (lower right) from the immunization experiment shown in (D). Each dot represents an individual mouse (n=2-5 mice per group). * p≤0.05; ** p≤0.005; **** p≤0.0001; two-tailed t-test.

To assess the activity of the Rosa26-AID allele we first performed *in vitro* stimulations of spleen naïve B cells in the presence of LPS and IL4 and found that the frequency of CSR to IgG1 was increased in R26^+/AID^ Cd19^+/cre^ B cells when compared to R26^+/+^ Cd19^+/cre^ control B cells (Figure 1C). We next assessed GC formation and CSR *in vivo* after mouse immunization with sheep red blood cells (SRBC). We found that R26^+/AID^ Cd19^+/cre^ mice accumulated more IgG1+ switched cells than did R26^+/+^ Cd19^+/cre^ cells, while the proportion of Fas+ GL7+ GC B cells was not affected (Figure 1D and 1E). We conclude that Rosa26^+/AID^ Cd19-Cre^ki/+^ mice display increased AID expression and activity.

### Early AID supra-expression is not sufficient to promote B cell lymphoma

To evaluate the contribution of AID supra-expression to B cell lymphomagenesis we generated cohorts of R26^+/+^ Cd19^+/cre^ and R26^+/AID^ Cd19^+/cre^ mice for ageing experiments. Mice were inspected weekly and animals showing symptoms of overt disease were euthanized. We found that survival was not affected in R26^+/AID^ Cd19^+/cre^ compared to R26^+/+^ Cd19^+/cre^ mice (Figure 2A, left) and that lymphoma incidence was not increased in R26^+/AID^ Cd19^+/cre^ mice (Figure 2A, right, and not shown). Cre-driven expression of AID in R26^+/AID^ Cd19^+/cre^ mice occurs progressively and relatively late during B cell differentiation. To assess whether the lack of oncogenic impact of AID in R26^+/AID^ Cd19^+/cre^ mice was due to a specific resistance of mature B cells to AID oncogenic transformation, we generated R26^+/AID^ Vav-Cre^+/TG^ mice, where AID supra-expression is driven by Vav-Cre from very early hematopoietic progenitors ^53^. Reporter GFP expression was detected from early lymphoid development as expected (Figure S2). We again found no survival reduction and no apparent histological differences in R26^+/AID^ Vav-Cre^+/TG^ mice compared to control R26^+/+^ Vav-Cre^+/TG^ mice (Figure 2B). Likewise, R26^+/AID^ mb1^+/cre^ mice did not show increased lymphoma incidence compared to R26^+/+^ mb1^+/cre^ littermates ^43^.

**Figure 2.**
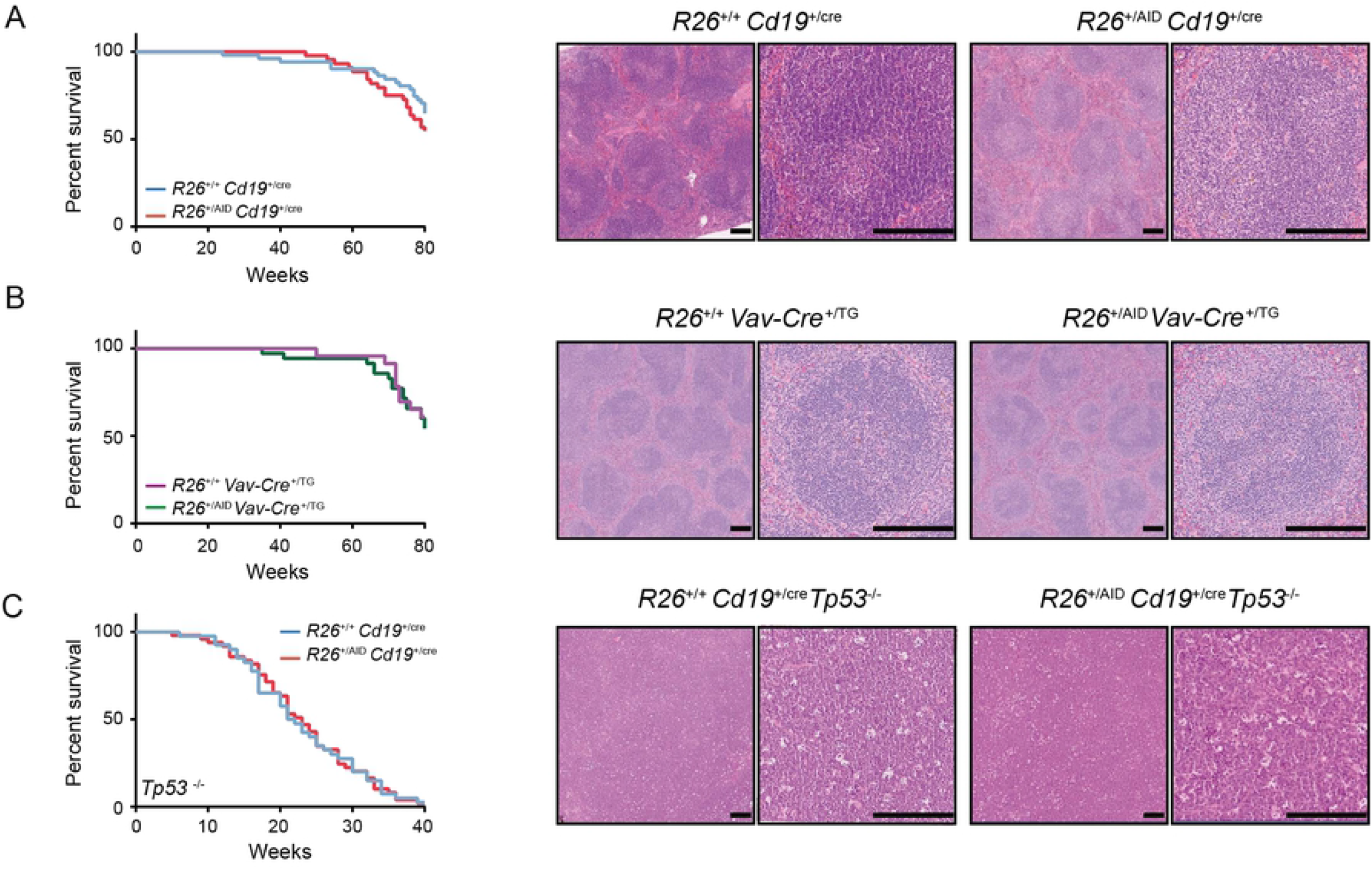
R26^+/AID^Cd19^+/Cre^ mice do not develop lymphoma. Kaplan-Meier survival analysis (left) and representative H&E staining of spleen sections (right) of **(A)** R26^+/+^Cd19^+/Cre^ (n=51) and R26^+/AID^Cd19^+/Cre^ mice (n=44) mice and **(B)** R26^+/+^ Vav ^+/cre^ (n=23) and R26^+/AID^ Vav ^+/cre^ (n=35) mice. **(C)** Kaplan-Meier survival analysis of R26^+/+^Cd19^+/Cre^Tp53^-/-^ (n=40) and R26^+/AID^Cd19^+/Cre^Tp53^-/-^ (n=49) and representative H&E staining of thymus T cell lymphomas found in both groups of mice. Mice were followed for 80 weeks (A-B) or at humane endpoint (C). p>0.05 in all cases; log-rank Mantel-Cox test. Magnification is 5x (left) and 20x (right); magnification bar is 200μm.

It has previously been shown in a different mouse model that deregulated expression of AID is insufficient to induce B cell lymphoma and that concomitant loss of p53 is required for transformation ^48^. To address whether in our R26^+/AID^ Cd19^+/cre^ mice p53 tumor suppressor activity was preventing AID-induced oncogenic transformation, we generated R26^+/AID^ Cd19^+/cre^ Tp53^-/-^ mice. We found that survival was not affected in R26^+/AID^ Cd19^+/cre^ Tp53^-/-^ mice when compared to R26^+/+^ Cd19^+/cre^ Tp53^-/-^ (Figure 2C, left). Likewise, the majority of tumor-bearing mice succumbed to T cell lymphoma or solid tumors both in R26^+/+^ Cd19^+/cre^ Tp53^-/-^ and in R26^+/AID^ Cd19^+/cre^ Tp53^-/-^ mice (65% and 77% combined frequency, respectively).

Thus, supra-physiological expression of AID in R26^+/AID^ mice did not impact on the overall mouse survival or the incidence of B cell lymphomas, regardless of the onset of expression or the absence of p53 tumor suppressor.

### UNG deficiency increases mutation load in B cells from R26^+/AID^ Cd19^+/cre^ mice

We have previously shown that the combined action of BER and MMR can faithfully repair a big fraction of the mutations triggered by AID deamination ^19,29^. To determine if under supra-physiological AID expression abolishing BER would result in a higher mutation load, we generated R26^+/AID^ Cd19^+/cre^ Ung^-/-^ mice. AID-induced SHM frequency was assessed at the Sμ region of the IgH locus by PCRseq ^29^ in naïve B cells or in GC B cells isolated from SRBC-immunized mice. As controls, we used naïve or GC B cells from R26^+/+^ Cd19^+/cre^ Ung^+/-^ mice, R26^+/AID^ Cd19^+/cre^ Ung^+/-^ mice and R26^+/+^ Cd19^+/cre^ Ung^-/-^ mice. In addition, we included LPS+IL4 activated B cells from AID deficient mice (Aicda^-/-^) as a negative control. We found that in naïve B cells, early AID supra-expression triggered detectable accumulation of mutations only in the absence of UNG (R26^+/AID^ Cd19^+/cre^ Ung^-/-^ versus R26^+/AID^ Cd19^+/cre^ Ung^+/-^or R26^+/+^ Cd19^+/cre^ Ung^-/-^) (Figure 3A, left). We obtained similar results in GC B cells isolated from SRBC-immunized mice, where mutation load driven by Rosa26-AID is very much increased when combined with UNG deficiency, but shows only as a trend in an UNG proficient background (Figure 3A, right). Separate analysis of transition and transversion mutations showed that increase in mutation frequency in UNG deficient cells was accounted by an increase in transitions at G/C pairs while no increase in transversion mutations was detected (Figure 3B), consistent with the outcome of direct replication of unrepaired U:G mismatches ^14,54^.

**Figure 3.**
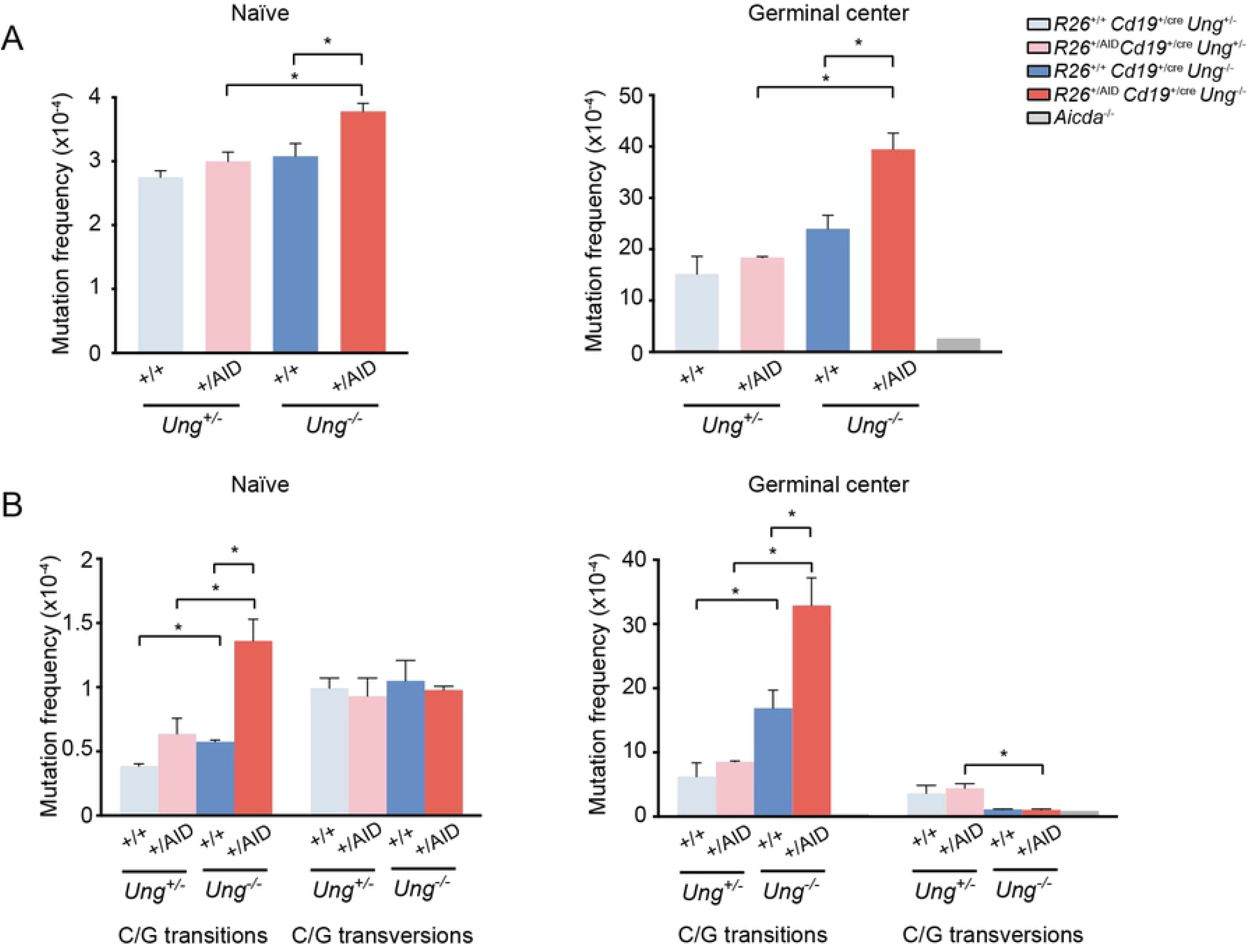
Increased SHM in BER deficient mice. Mutation analysis of IgH Sμ region in naïve and GC B cells from R26^+/+^ Cd19^+/cre^ Ung^+/-^, R26^+/AID^ Cd19^+/cre^ Ung^+/-^, R26^+/+^ Cd19^+/cre^ Ung^-/-^, R26^+/AID^ Cd19^+/cre^ Ung^-/-^ and Aicda^-/-^ mice by PCR-Seq. (A) Total mutation frequency and (B) frequency of transition and transversion mutations at C/G lying in WRCY/RGYW hotspots. (n=2, each composed of a pool of 3 mice; * p ≤ 0.05; two-tailed t-test; error bars represent STD).

We conclude that UNG deficiency synergistically increases the mutation load in B cells that supra-express AID. Given that chromosome translocations are impaired in UNG deficient B cells^24^, R26^+/AID^ Cd19^+/cre^ Ung^-/-^ mice provide a unique opportunity to assess the contribution of AID-induced SHM to lymphomagenesis.

### UNG deficiency promotes oncogenic transformation by AID

To address the contribution of AID-induced SHM to oncogenic transformation, we generated cohorts of R26^+/+^ Cd19^+/cre^ Ung^+/-^ mice, R26^+/AID^ Cd19^+/cre^ Ung^+/-^ mice and R26^+/+^ Cd19^+/cre^ Ung^-/-^ and R26^+/AID^ Cd19^+/cre^ Ung^-/-^ mice and performed ageing experiments. We found that R26^+/AID^ Cd19^+/cre^ Ung^-/-^ mice showed a reduced median survival compared to the other groups of mice (Figure 4A). Moreover, at sacrifice R26^+/AID^ Cd19^+/cre^ Ung^-/-^ mice harboured a different pattern of B cell lymphomas by immuhistochemistry evaluation, with a smaller proportion of large-B cell lymphoma (DLBCL-like) and a bigger proportion of small B cell lymphoma (FL-like) than R26^+/+^ Cd19^+/cre^ Ung^-/-^ mice (Figure 4B-D). Tumors retained the expression of AID and typically expressed B cell markers, including Pax or B220 (Figure 4C-D) as well as the proliferation antigen Ki67. In addition, we observed that the majority of the tumors expressed GL7 and Bcl-6 (Figure 4C-D and Supplementary Figure S3), supporting a GC origin. Together, these results indicate that under the condition of UNG deficiency, AID supra-expression promotes GC B cell transformation.

**Figure 4.**
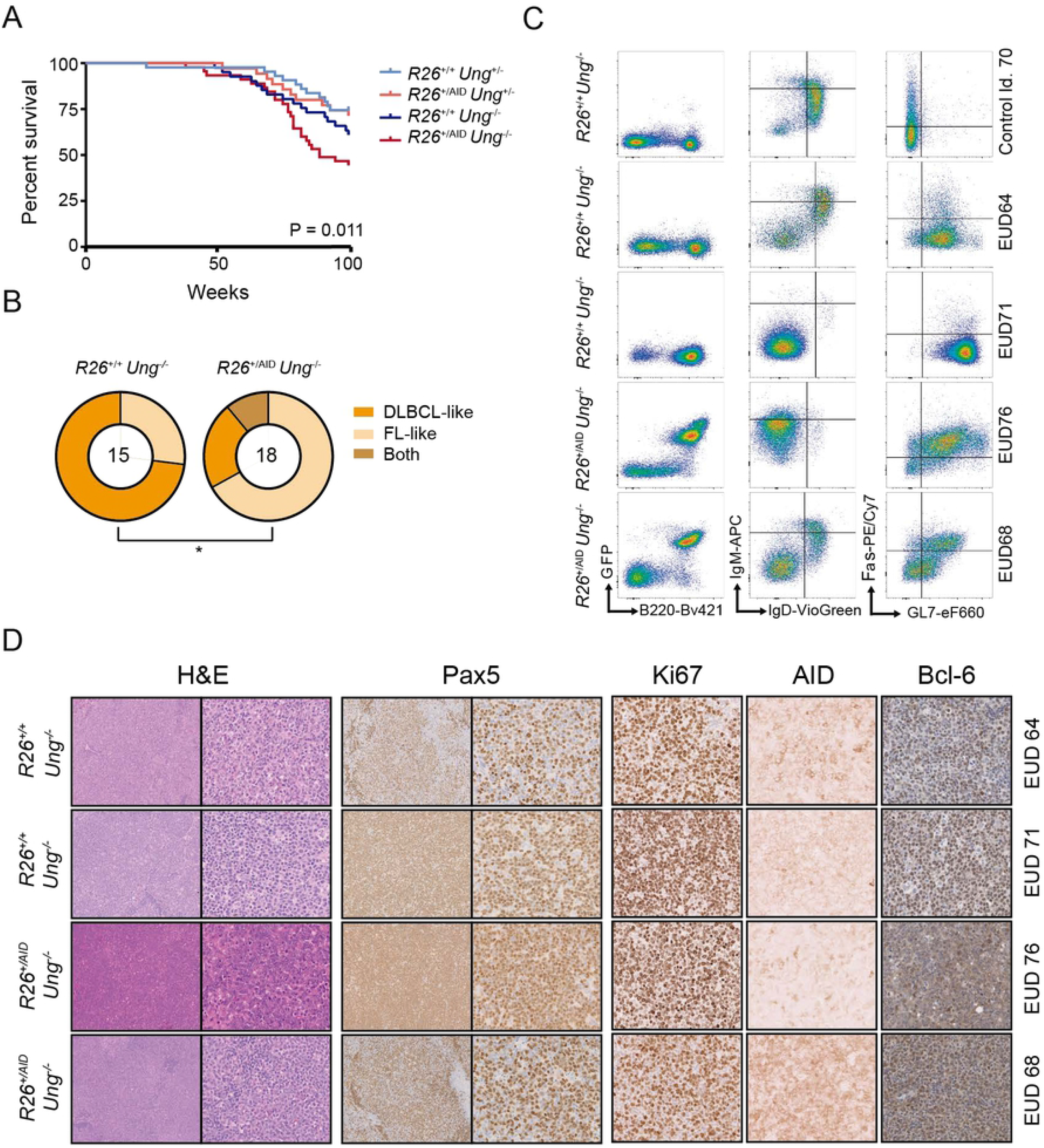
BER deficiency promotes AID-dependent lymphomagenesis. **(A)** Kaplan-Meier survival analysis of R26^+/+^Cd19^+/cre^Ung^+/-^(n=43), R26^+/AID^Cd19^+/cre^Ung^+/-^ (n=35), R26^+/+^Cd19^+/cre^Ung^-/-^ (n=41) and R26^+/AID^Cd19^+/cre^Ung^-/-^ (n=45) mice (p=0.011, Log-rank Mantel-Cox test). **(B)** Lymphoma diagnosis of R26^+/+^Cd19^+/cre^Ung^-/-^ and R26^+/AID^Cd19^+/cre^Ung^-/-^ mice. Outer rings depict proportion of lymphomas with a FL-like or DLBCL-like phenotype; inner circle indicates the number of lymphoma tumors analyzed. * p ≤ 0.05, Two-tailed Fisher test. **(C)** Immunophenotype of a control mouse spleen (upper panels) and of representative tumors from R26^+/+^Cd19^+/cre^Ung^-/-^ and R26^+/AID^ Cd19^+/cre^ Ung^-/-^ mice (lower panels). Total live cells (left panel) and B220+ gated cells (middle and right panels) are shown. **(D)** Immunohistochemistry analysis of representative tumors from R26^+/+^Cd19^+/cre^Ung^-/-^ and R26^+/AID^Cd19^+/cre^Ung^-/-^ mice. Magnification is 10x for H&E and Pax5 (left panels) and 40x for the rest of the panels.

### AID promotes intratumor heterogeneity

To analyse the contribution of AID to lymphomagenesis at the molecular level, we performed Whole Exome Sequencing (WES) of R26^+/+^ Cd19^+/cre^ Ung^-/-^ (n=7) and R26^+/AID^ Cd19^+/cre^ Ung^-/-^ (n=8) tumors and a healthy tissue control. We sequenced a total of 148Gb with an average depth of 79x (Table S1, Fig. S4). Variant calling analysis of tumoral versus healthy tissue (see Methods for details) revealed a total number of 4801 unique variants in 5195 different genes, with an average of ~640 variants and ~740 genes affected per tumor (Fig. S4A). Most of the variants corresponded to single nucleotide variants (SNVs, ~76%) and a minor fraction were insertions or deletions (INDELS) (~24%) (Fig. S4A, B).

To assess the mutation load of R26^+/+^ Cd19^+/cre^ Ung^-/-^ and R26^+/AID^ Cd19^+/cre^ Ung^-/-^ tumors, we first calculated mutation frequency per genotype. For this analysis (Figure 5A) only transition mutations at G/C pairs were considered. This approach allowed us to identify very large numbers of mutations at C/G pairs, which were preferentially enriched within WRC/GYW (W = A/T; R = C/G; Y = C/T) AID mutational hotspots (Figure 5A). Interestingly, we found a significantly higher mutation frequency and more pronounced hotspot focusing in R26^+/AID^ Cd19^+/cre^ Ung^-/-^ than in R26^+/+^ Cd19^+/cre^ Ung^-/-^ tumors (Fig. 5A). These results indicate that tumors harbor AID-induced mutations and that they occur at larger numbers in R26^+/AID^ Cd19^+/cre^ Ung^-/-^ than in R26^+/+^ Cd19^+/cre^ Ung^-/-^ tumors, consistently with our findings in non-transformed GC B cells. We found that copy number variations occurred at similar frequencies in R26^+/AID^ Cd19^+/cre^ Ung^-/-^ and R26^+/+^ Cd19^+/cre^ Ung^-/-^ tumors (Fig. S4C), in agreement with the idea that AID activity is not expected to give rise to double strand breaks in the absence of UNG ^10,14,24,55^.

**Figure 5.**
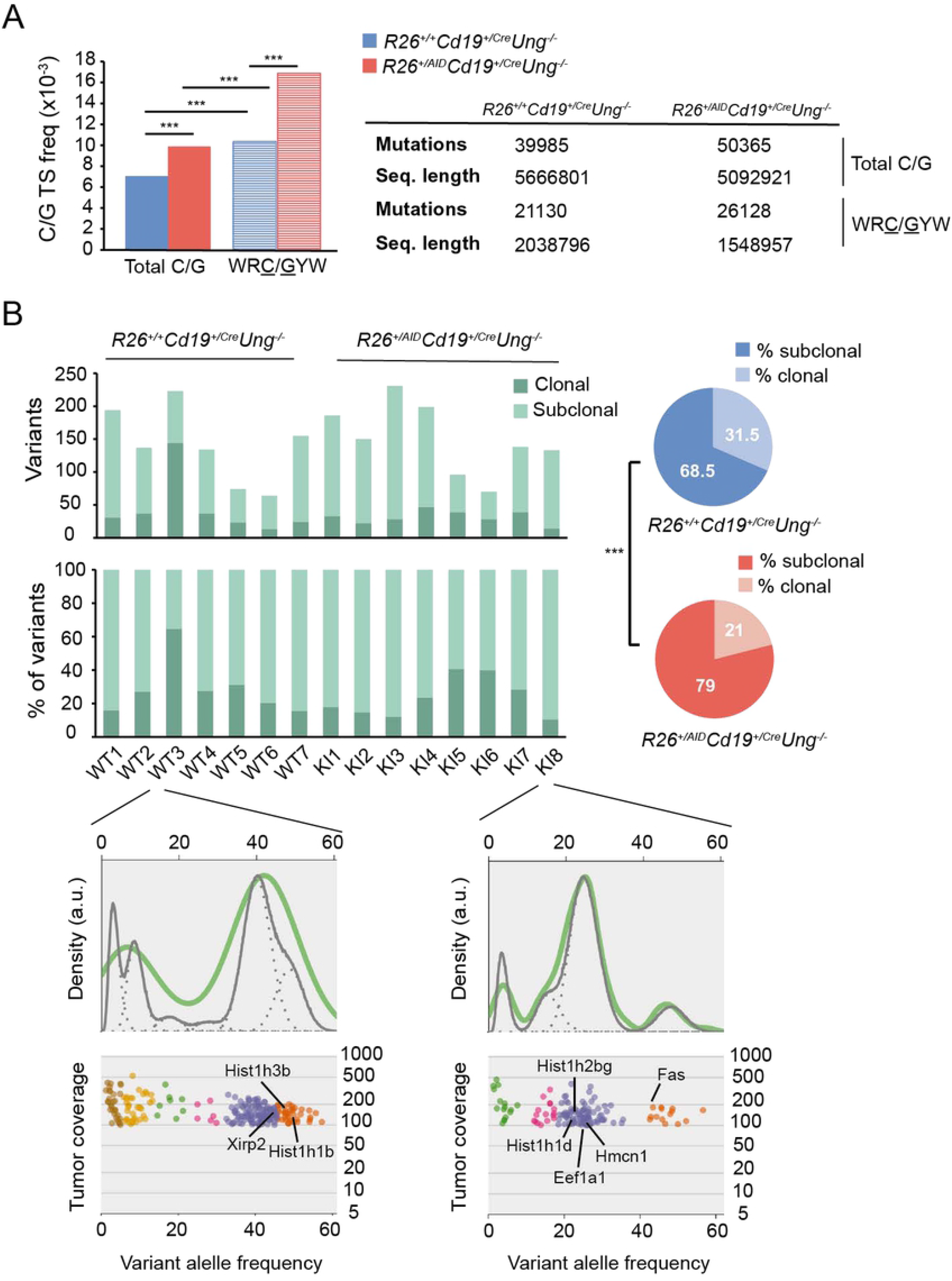
AID contributes to intratumor heterogeneity. **(A)** Average transition mutation frequency at total C/G pairs and at C/G within WRC/GYW hotspots in tumors from R26^+/+^Cd19^+/cre^Ung^-/-^ and R26^+/AID^Cd19^+/cre^Ung^-/-^ mice as measured by whole exome sequencing (Two-tailed γ2 test; ***p ≤10^-3^). **(B)** Number (upper panel) and proportion (lower panel) of clonal and subclonal variants in each of the seven R26^+/+^Cd19^+/cre^Ung^-/-^ (WT) and eight R26^+/AID^Cd19^+/cre^Ung^-/-^ (KI) tumors analyzed. Piecharts depict the proportion of clonal and subclonal variants of all the tumors analyzed pooled by genotype (Two-tailed Fisher test; ***p ≤10^-3^). Inset panels show a representative example of the intratumor heterogeneity in each genotype. Clusters as defined by SciClone are depicted in different colors. Clonal variants are those belonging to the cluster with the highest variant allele frequency (VAF; orange clusters in inset panels), usually between 40-50%. Subclonal variants are those belonging to clusters with a lower VAF than that of the clonal cluster. Density plots depict the posterior predictive density, a surrogate of the accuracy of the prediction model assigning variants to the different clusters (green lines represent the diploid model; grey continuous and discontinuous lines represent the model fit of each cluster and each variant, respectively). Tumor coverage refers to the number of reads supporting each variant. Genes that are frequently mutated in human lymphoma tumors are indicated in black.

Intratumor heterogeneity (ITH) has been related to poor prognosis and impaired response to treatment ^56–58^. To assess if AID-induced SHM could contribute to ITH we measured clonal and subclonal variants in R26^+/+^ Cd19^+/cre^ Ung^-/-^ and R26^+/AID^ Cd19^+/cre^ Ung^-/-^ tumors as a proxy to estimate ITH ^59^. In this context, clonal variants refer to early acquired mutations shared by a big proportion of the cancer cells, as reflected by their high variant allele frequency (VAF). Interestingly, several genes that are frequently mutated in human lymphomas, such as *Fas^60^, Eeflal^61^* and some members of the histone family genes (*Hist1h1b, Hist1h1d, Hist1h2bg, Hist1h3b*)^62^ bear clonal variants (Fig. 5B, inset). Conversely, subclonal variants are mutations arising later in tumor development, defining younger clones with lower VAFs. Thus, a higher proportion of subclonal variants in a tumor reflects a higher ITH. We found that R26^+/AID^ Cd19^+/cre^ Ung^-/-^ tumors have a higher proportion of subclonal variants than R26^+/+^ Cd19^+/cre^ Ung^-/-^ tumors (Figure 5B, right), indicating that AID contributes to the generation of ITH.

To gain insights into the functional relevance of AID activity to the development of lymphoma, we focused on variants affecting known cancer-related genes, including genes that are frequently mutated in human lymphoma ^63–69^ see Methods for details. We identified a total number of 132 and 154 somatically mutated cancer-related genes (see Methods) in R26^+/+^ Cd19^+/cre^ Ung^-/-^ and R26^+/AID^ Cd19^+/cre^ Ung^-/-^ tumors, respectively (Table S2; complete list in Table S3). Notably, a big proportion of which are shared between the two groups of tumors (Figure 6A), suggesting a common network of functional pathways being affected for lymphoma generation (Table S3; Figure 6B). However, we identified several cancer-related genes that were exclusively mutated in R26^+/AID^ Cd19^+/cre^ Ung^-/-^ tumors (Table S3; Figure 6A). Functional annotation revealed that cancer-related genes mutated in R26^+/AID^ Cd19^+/cre^ Ung^-/-^ tumors impinge on more diverse cellular functions, with the p53 DNA damage checkpoint or the negative regulation of B cell activation as well as cell proliferation or PI3K signalling being more affected in R26^+/AID^ Cd19^+/cre^ Ung^-/-^ than in R26^+/+^ Cd19^+/cre^ Ung^-/-^ tumors (Figure 6B, C). Furthermore, mutations in R26^+/AID^ Cd19^+/cre^ Ung^-/-^ exclusive cancer genes occurred more frequently in C/G pairs than in A/T pairs and were enriched in hotspots, strongly suggesting that they have originated from AID activity (Figure 6D). Together, these data indicate that AID induced SHM increases ITH and broadens the range of oncogenic events associated to lymphomagenesis.

**Figure 6.**
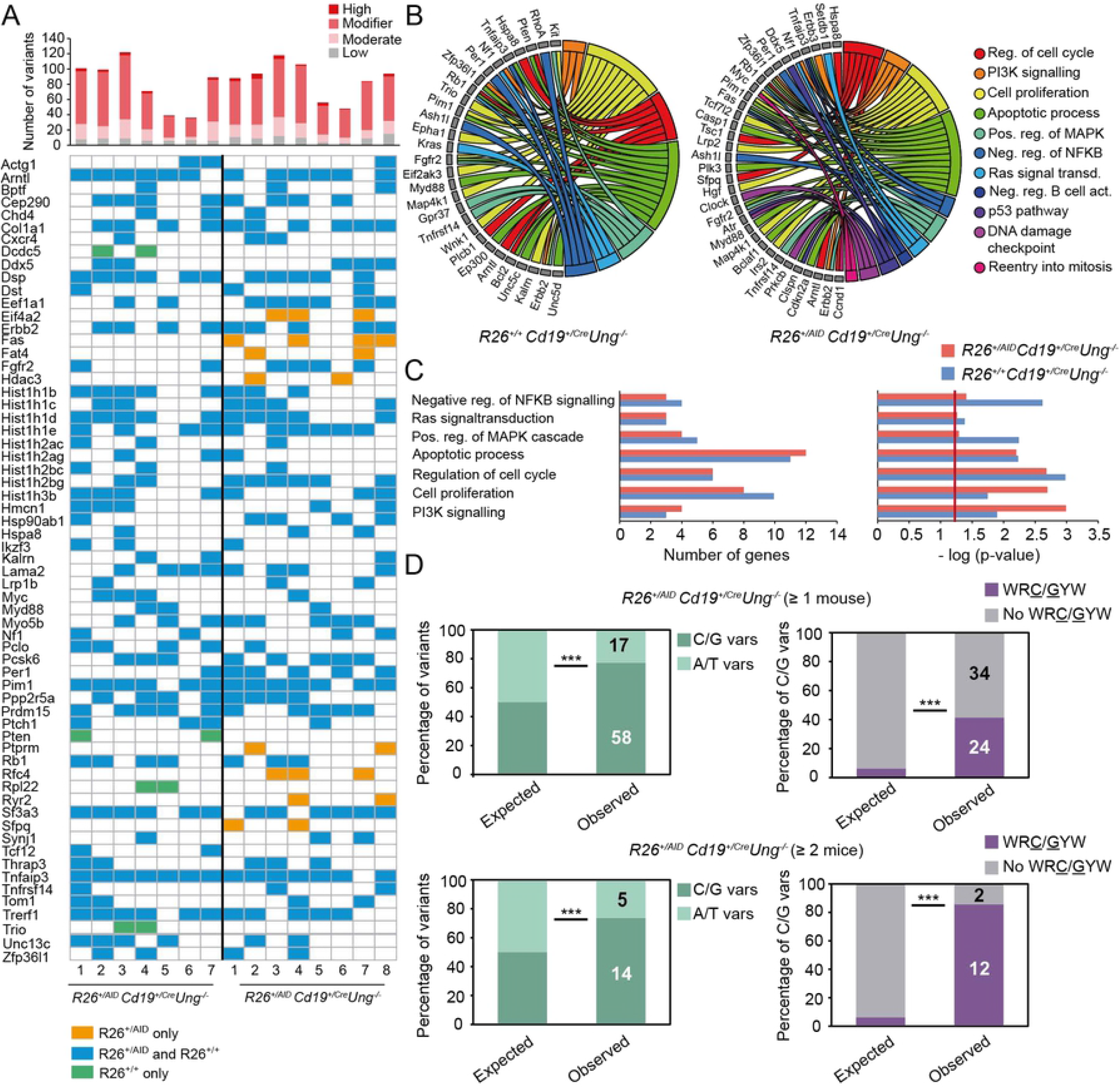
AID broadens the functional impact of oncogenic events in Ung deficient tumors. (A) Cancer-related genes found mutated in tumors from R26^+/+^ Cd19^+/cre^ Ung^-/-^ and R26^+/AID^ Cd19^+/cre^ Ung^-/-^ mice. Upper panel shows total number of variants identified in all cancer-related genes in each of the tumors; colors code defines the functional impact of the variants as annotated by Variant Effect Predictor (VEF; see methods section for details). Lower panel represents presence (colored box) or absence (empty box) of variants in each of the cancer-related genes for each of the analyzed tumors. For simplicity, only mutated genes found in >= 2 tumors per group are shown; a full list is included in Table S3. Green indicates genes exclusively mutated in R26^+/+^Cd19^+/cre^Ung^-/-^ tumors; orange indicates genes exclusively mutated in R26^+/AID^Cd19^+/cre^Ung^-/-^ tumors; blue indicates genes mutated in both. **(B)** Circos plot representation of the biological functions associated to all mutated cancer-related genes (Gene Ontology biological process database) in R26^+/+^Cd19^+/cre^Ung^-/-^ (left) and R26^+/AID^Cd19^+/cre^Ung^-/-^ (right) tumors. **(C)** Number of genes (left) and p-value of the enrichment (right, EASE score-modified Fisher exact test-as provided by DAVID, see methods for details) of the common biological functions affected in both R26^+/+^Cd19^+/cre^Ung^-/-^ and R26^+/AID^Cd19^+/cre^Ung^-/-^ tumors. **(D)** Proportion of mutations in C/G and A/T nucleotides (left) and within WR<u>C</u>/<u>G</u>YW hotspots (right) in cancer-related genes that are exclusively mutated in R26^+/AID^Cd19^+/cre^Ung^-/-^ tumors (Fisher two-tailed test; ***p≤10-3).

## DISCUSSION

Here we have made use of a genetic model for AID expression to assess the contribution of AID to lymphoma development. Our R26^+/AID^ Cd19^+/cre^ model is different from previous approaches in that it is not a transgenic model and it allows conditional expression of AID together with GFP as a positive tracer for AID expression. Expectedly, we find that AID mRNA and protein are detected in naïve B cells in R26^+/AID^ Cd19^+/cre^ and that in activated B cells total AID expression and activity is increased in R26^+/AID^ Cd19^+/cre^ mice compared to R26^+/+^ Cd19^+/cre^ littermates. Thus, in contrast to previous reports ^45–47^, heterologous Rosa26-AID expression in R26^+/AID^ Cd19^+/cre^ mice does not seem subject to particular mechanisms of negative regulation, and simply leads to a relatively proportional increase of AID activity.

In agreement with previous data using different mouse models ^44–48^, we show here that AID supra-expression in R26^+/AID^ mice is not sufficient to promote mature B cell transformation even if driven from very early on in B cell progenitors with Vav-Cre or mb1-Cre ^43^ drivers, suggesting that cumulative AID activity during B cell differentiation does not have a measurable oncogenic impact by itself. Interestingly, Rosa26-AID did not promote mature B cell lymphomas even in the absence of p53, in sharp contrast with a previous report ^48^. This discrepancy may well be explained by the difference in AID expression levels achieved in these two studies. While AID expression and activity was only mildly increased in R26^+/AID^ Cd19^+/cre^ mice, it reached much higher levels in the IgκAID mouse from Robbiani et al, when compared to wild type mice, specially in GC B cells. We believe that these expression levels may promote increased genome instability and hence triggered more efficiently a DNA damage response, which would explain the increased incidence of B cell lymphoma observed in the absence of p53 ^48^.

In contrast with the lack of lymphomagenic effect of AID, alone or combined with p53 deficiency, we show here that in the absence of UNG, a mild increase in AID expression (R26^+/AID^ Cd19^+/cre^ mice) is sufficient to decrease mean survival and contributes to lymphoma evolution. This in turn can be a direct reflect of the increased SHM frequency observed when UNG deficiency is combined with Rosa26-AID expression, while Rosa26-AID alone (this study) or Ung^-/-^ alone (this study, ^19,29^) have only a moderate effect on SHM. Given that UNG is essential for AID-triggered double strand breaks and chromosome translocations, our report provides the first direct genetic proof that the SHM activity of AID can, by itself, contribute to B cell transformation. In turn, the absence of double strand breaks and thus of DNA damage response could favour the survival and accumulation of mutations of GC B cells and their likelihood to oncogenic transformation. Importantly, mutations and homozygous UNG losses have been found in human B-NHL^70^.

Various studies have previously suggested that AID can drive or contribute to clonal evolution in B cell malignancies ^35,71,72^. Our study also provides, to our knowledge, the first causal evidence of AID-induced SHM B cell tumors, as measured by whole exome sequencing in a genetic model. Interestingly, both control R26^+/+^ Cd19^+/cre^ and R26^+/AID^ Cd19^+/cre^ tumors bear the hallmark of AID activity, but this is more pronounced in tumors with supra-expression of AID (R26^+/AID^ Cd19^+/cre^ mice). Further, our study directly shows that AID-induced SHM promotes a bigger proportion of subclones and increased ITH. Our model predicts that AID would increase ITH and contribute to second hit oncogenic transformation events preferentially in tumoral cells with UNG loss of function events. Thus, we propose that AID-induced SHM may contribute to lymphoma progression by giving rise to a more diverse output of oncogenic variants that presumably impinge on tumor evolution and aggressiveness ^56–58^.

## METHODS

### Mice and B cell cultures

R26^+/AID^ mice were described previously ^49^ and bred to Cd19^cre 50^, Vav-Cre^TG/+ 53^ *Tp53*^-/-^ ^73^ and Ung^-/- 74^ mice as indicated. *Aicda*^-/-9^ were used as negative control for SHM assays. Littermates aged from 6 to 12 weeks were used for SHM, CSR and immunization experiments. All animals were housed in the Centro Nacional de Investigaciones Cardiovasculares animal facility under specific pathogen-free conditions. All animal procedures conformed to EU Directive 2010/63EU and Recommendation 2007/526/EC regarding the protection of animals used for experimental and other scientific purposes, enforced in Spanish law under RD 53/2013. The procedures have been reviewed by the Institutional Animal Care and Use Committee (IACUC) of Centro Nacional de Investigaciones Cardiovasculares, and approved by Consejeria de Medio Ambiente, Administración Local y Ordenación del Territorio of Comunidad de Madrid.

T-dependent immunization was induced by intravenously injection of 10^8^ sheep red blood cells resuspended in 100 μl of sterile PBS. Immunization response was analyzed in spleen 7 days after sheep red blood cell injection.

For CSR experiments spleen naive B cells were isolated by immunomagnetic depletion using anti-CD43 beads (Miltenyi Biotec). Purified naïve B cells were cultured at 1.2 x 10^6^ cells/ml in complete RPMI medium supplemented with 10% FBS (Gibco), 50 mM 2-βmercaptoethanol (Gibco), 10 mM Hepes (Gibco), penicillin and streptomycin (Gibco), 25 μg/ml lipopolysacharide (LPS, Sigma-Aldrich) and 10 ng/ml IL-4 (PreproTech).

### Flow cytometry and cell sorting

Single-cell suspensions were obtained from bone marrow, spleen or cultured B cells and stained with fluorochrome/biotin-conjugated antibodies to detect mouse surface antigens: B220 (RA3-6B2), CD19 (1D3), IgM (ll/41), IgD (11-26c.2a), IgG1 (A85-1), CD25 (PC61), Fas (Jo2)), Ly77 (GL7), CD4 (RM4-5), CD8 (53-6.7) and CD44 (IM7). Samples were acquired on LSRFortessa or FACSCanto instruments (BD Biosciences) and analyzed with FlowJo software. GC B cells and naïve B cells were purified from spleen from immunized mice using FACSAria (BD Biosciences) or SY3200 (Sony).

### qRT-PCR

RNA was extracted from naïve and cultured B cells using Trizol and RNeasy kit and Dnase treatment (Qiagen) and retro-transcribed to cDNA using random hexamers (Roche) and SuperScript II reverse transcriptase (Invitrogen). qRT-PCR was performed using Sybr Green in the 7900 Fast Real Time thermocycler (Applied Biosystems). The following primers were used: mouse-GAPDH (forward) 5’-TGA AGC AGG CAT CTG AGG G-3’, (reverse) 5’-CGA AGG TGG AAG AGT GGG AG-3’; Total AID (forward) 5’-ACC TTC GCA ACA AGT CTG GCT −3’, (reverse) 5’-AGC CTT GCG GTC TTC ACA GAA −3’ and R26-AID (forward) 5’-ATT CGC CCT TGG GAT CCA-3’, (reverse) 5’-AGC CAG ACT TGT TGC GAA GGT-3’. mRNA expression was normalized to GAPDH mRNA measured in triplicates in each sample. Relative quantification was performed using the ΔΔC_T_ method.

### Immunoblotting

Naïve and activated B cells were incubated on ice for 20 min in NP-40 lysis buffer in the presence of protease inhibitors (Roche) and lysates were cleared by centrifugation. Total protein was size-fractionated on SDS-PAGE 12% acrylamide-bisacrylamide gels and transferred to Protan nitrocellulose membrane (Whatman) in transfer buffer (0.19M glycine, 25 mM Tris base and 0.01% SDS) containing 20% methanol (90 min at 0.4A). Membranes were probed with anti-mouse-AID (1/25, eBioscience, 14-5959-82) and anti-mouse-tubulin (1/5000, Sigma-Aldrich, T9026). Then, membranes were incubated with HRP-conjugated anti-rat (1/5000, Bethyl Laboratories, A110-105P) and antimouse (1/10000, DAKO) antibodies, respectively and developed with Clarity™ Western ECL Substrate (Bio-Rad).

### IgH Sμ amplification and mutation analysis by PCR-Seq

For the analysis of mutations at IgH Sμ region, genomic DNA was isolated from GC and naïve B cells from immunized mice. Amplifications were performed in 4 independent reactions using the following primers:

(forward) 5’-AATGGATACCTCAGTGGTTTTTAATGGTGGGTTTA-3’; (reverse) 5’-GCGGCCCGGCTCATTCCAGTTCATTACAG-3’. Amplification reactions were carried in a final volume of 25 μl each, using 2.5 U Pfu Ultra HF DNA polymerase (Agilent) for 26 cycles (94°C for 30”, 55°C for 30”, 72°C for 60”). PCR products were purified and fragmented using a sonicator (Covaris), and libraries were prepared according to the manufacturer’s instructions (NEBNext Ultra II DNA Library Prep; New England Biolabs). Sequencing was performed in a HiSeq 2500 platform (Illumina). Analysis was performed as previously described ^29^.

### Immunohistochemistry

Spleens and tumors were fixed with 10% neutral-buffered formalin and embedded in paraffin. 5 μM sections were stained with hematoxylin and eosin (Sigma) using standard protocols. For Ki67, Bcl6 and Pax5 immunohistochemistry, sodium citrate buffer (10mM pH 6, Sigma-Aldrich) was used for antigen retrieval. Endogenous peroxidases were blocked with 3% (v/v) H_2_O_2_ (Merck) in methanol (Sigma-Aldrich). For AID immunohistochemistry, EDTA buffer (1mM EDTA, 0,05% Tween-20 pH 8) was used for antigen retrieval and endogenous peroxidases were blocked with 0.3% H_2_O_2_ in water. Rabbit anti-Ki67 (Abcam, 1/100), goat anti-Pax5 (Santa Cruz Biotechnology, 1/2000), rat anti-AID (eBioscience, 1/50), and anti-Bcl-6 (IG91E, generously provided by Dr G Roncador, CNIO) were incubated overnight at 4°C. Biotinylated secondary antibodies were incubated 1 hour at room temperature. Biotinylated antibodies were detected with the ABC system (Vector Laboratories). Images were acquired with a Leica DM2500 microscope or digitized in a Hamamatsu NanoZoomer Scanner. Tumors were diagnosed by a pathologist.

### Whole Exome Sequencing (WES)

Exome sequencing was performed in 15 tumors and one healthy tissue (tail) as a non-tumoral control. In brief, genomic DNA was isolated by standard procedures, quantified in a fluorometer (Qubit; Invitrogen), fragmented in a sonicator (Covaris) to ~200 nucleotide-long (mean size) fragments and purified using AMPure XP beads (Agencourt). Quality was assessed with the 2100 Bioanalyzer (Agilent). Then, fragment ends were repaired, adapters were ligated, and the resulting library was amplified and hybridized with a SureSelect Mouse All Exon kit library (Agilent) of RNA probes. DNA–RNA hybrids were then captured by magnetic bead selection. After indexing, libraries were paired-end sequenced in a HiSeq 2500 platform to produce 2×100bp reads. DNA capture, library preparation, and DNA sequencing were performed by the Genomics Unit at Centro Nacional de Investigaciones Cardiovasculares (CNIC).

#### a. Identification of somatic variants from WES data

Raw reads were demultiplexed and processed into *fastq* files by CASAVA (v1.8) and quality checked by FastQC (0.10.1). Sequencing adaptors were removed by cutadapt (1.7.1) (minlength=30; minoverlap=7) and the resulting reads were mapped to the mouse genome (GRCm38 v75 Feb 2014) using BWA-MEM (0.7.10-r789) (command line option -M to mark shorter split hits as secondary). Duplicates were marked by Picard tools (1.97). Variant calling was performed by means of GenomeAnalysisTK-3.5 (GATK3.5) and GATK Resource Bundle GRCm38 using default parameters and thresholds: 1) Aligned reads were realigned for known insertion/deletion events (Realigner TargetCreator and IndelRealigner tools) to minimize the number of mismatching bases and reduce false SNV calls; 2) base qualities were recalibrated (BaseRecalibrator tool) by a machine learning model to detect and correct systematic errors in base quality scores; 3) a custom Panel Of Normals (PON) was built; 4) somatic variants were called by MuTect2 (from GATK3.6); 5) Variants were functionally annotated with Variant Effect Predictor (VEP) version 84 (command line option --per-gene to output only the most severe functional consequence of the variant at the gene level) and filtered out to remove false positives (custom scripting). PON was built by combining the following variants: Set1, intersection between germline (HaplotypeCaller) and somatic (MuTect2) variant calls in healthy tissue; set2, somatic variants (MuTect2) that have been called in all the 15 tumor samples and healthy tissue, including variants labeled as “NO PASS” by the caller in ≤2 of the samples; set3, somatic variants (MuTect2) that have been called in ≥13 tumor samples and healthy tissue and that have been labeled as “PASS” by the caller in all the samples. Filtering out of potential false positives was done by removing all variants that were called in healthy tissue, annotated in dbSNP (v142) or annotated in the Mouse Genomes Project (v5) for 129P2/OlaHsd, 129S1/SvImJ, 129S5SvEvBrd, BALB/cJ, C57BL/6NJ and CBA/J strains. Exome sequencing metrics included in TableS1 were calculated using Picard tools HsMetrics.

#### b. Copy number variation analysis from WES data

Copy number variation analysis was done using CopywriteR, an R package which implements a statistical method to estimate CNV from off-target reads in capture-based sequencing experiments. This approach overcomes most of the limitations and biases associated to CNV estimation from targeted sequencing data. Briefly, the software reads the alignment files, removes non-random off-target reads, calculates depth of coverage in predefined bins, corrects it by GC-content and mappability and generates a CNV profile. We used 20Kb bins and provided both tumor and healthy tissue (as a negative control) samples for the software to call CNV regions that exclusively occur in tumors.

#### c. Inference of clonal architecture of tumors and estimation of intratumor heterogeneity from WES data

Intratumoral heterogeneity was estimated using the R package SciClone ^59^. Variant allele frequencies (VAFs) and CNV information were used to infer clonal architecture of each of the 15 tumors analyzed. To ensure statistical robustness, only those variants supported by ≥ 100 reads were considered for the ITH analysis, as recommended by the authors of the tool. Mutations with a VAF greater than 0.8 were excluded to filter out germline mutations. Variants belonging to the cluster with the highest VAF were considered as “clonal” and those belonging to clusters with a lower VAF were annotated as “subclonal”.

#### d. Identification and functional annotation of cancer associated genes from WES data

Cancer associated genes were identified as those genes that bear at least one variant in one of the 15 tumors analyzed and that are annotated in IntOGen ^63^ as cancer drivers (gene set composed of 459 genes) or appear mutated in human diffuse large B cell lymphoma, Burkitt lymphoma or follicular lymphoma tumors ^63–69^ (lymphoma associated genes; gene set composed of 239 genes). Functional annotation of the 132 and 154 cancer-related genes identified in R26^+/+^ Cd19^Cre/+^ Ung^-/-^ and R26^+/ki^ Cd19^Cre/+^ Ung^-/-^ tumors was performed by Gene Ontology Biological process database. Enrichment tests were done using DAVID 6.8.

### Statistical analysis

Statistical analyses were performed with GraphPad version 6.01 or stats R package v3.1.1. Error bars in figures represent SD. Two-tailed Student’s t test was applied to continuous data following a normal distribution (Shapiro-Wilk test) and two-tailed Fisher or χ^2^ test were used to assess differences between categorical variables. Kaplan-Meier survival curves were compared by log-rank Mantel-Cox test. Differences were considered statistically significant at P ≤ 0.05.

## ACKOWLEDGEMENTS

We thank all members of the B cell biology lab for scientific discussion, JM Ligos and the CNIC Cellomics Unit for flow cytometry assistance, CNIC Genomics Unit for assistance with whole genome sequencing and PCRseq experiments, CNIC Histology Unit for assistance with immunohistochemistry stainings and Carlos Torroja for helpful advice with whole exome sequencing data analysis.

PD was supported by the AECC foundation (AIO2012). EM-Z is an FPI fellow (BES-2014-069525). AFA-P, S.M.M. and A.R.R. are supported by CNIC funding. This project was funded by the Spanish Ministerio de Economía, Industria y Competitividad SAF2013-42767-R, SAF2016-75511-R, and European Research Council StG BCLYM, to A.R.R. The CNIC is supported by the Ministerio de Ciencia, Innovacion y Universidades and the Pro-CNIC Foundation, and is a Severo Ochoa Center of Excellence (SEV-2015-0505).

## AUTHOR CONTRIBUTIONS

PD, AFA-P, EM-Z, IVS, SMM, LB and VGdY designed and performed experiments and analysed data. JdlB and FS-C performed whole exome sequencing analysis, MC and AdM did pathology evaluations. PD, AFA-P, EM-Z and VGdY wrote manuscript. ARR designed strategy and experiments and wrote manuscript.

## MATERIALS AND CORRESPONDENCE

Correspondence and material requests should be addressed to

